# Genetic erosion and projected habitat loss in the protected Alpine moth *Actias isabellae galliaegloria* (Lepidoptera: Saturniidae)

**DOI:** 10.64898/2026.02.19.706775

**Authors:** Flora Lambert-Auger, Marina Querejeta, Stéphane Boyer, Marta Vila, Carlos Lopez-Vaamonde, Jérémy Gauthier

## Abstract

1. Montane and specialist insects are particularly vulnerable to habitat loss and climate change. The Spanish Moon Moth (*Actias isabellae*), a rare and protected species, hosts an isolated Alpine population (subspecies *galliaegloria*) whose conservation status remains unclear.
2. We combined Restriction-site Associated DNA sequencing (RADseq) and Ecological Niche Modelling (ENM) to assess its genetic diversity, population structure, and environmental vulnerability.
3. Genomic analyses revealed extremely low genetic diversity and inbreeding in the Alpine subspecies, consistent with a historical bottleneck, reducing adaptive potential.
4. Species distribution models predict a major contraction of suitable habitat by 2050 due to rising temperatures and declines in its primary host, Scots pine (*Pinus sylvestris*).
5. These findings highlight the compounded risks posed by genetic impoverishment and habitat loss, emphasizing the urgent need for targeted conservation measures.
6. Our study demonstrates the value of integrating genomic and ecological approaches to evaluate extinction risk and guide management strategies for montane specialist insects under rapid environmental change.

## Introduction

Global biodiversity is declining at an unprecedented rate, with insects appearing to be particularly affected (Díaz *et al*., 2019; Warren *et al*., 2021; Edwards *et al*., 2025). Increasing evidence highlights climate change, habitat loss and agricultural activities as the major drivers of insect decline worldwide (Dirzo *et al*., 2014; Hallmann *et al*., 2017; Wagner *et al*., 2021). Habitat loss occurs both directly, through agriculture intensification, urbanisation and land-use changes, and indirectly, as climate change alters environmental conditions, species interactions and community composition. These pressures lead to population declines, local extinctions and fragmentation of remaining populations, increasing their vulnerability to stochastic events and accelerating biodiversity loss.

Climate change exacerbates these threats by shifting suitable habitats poleward and upward in elevation, disproportionally affecting cold-adapted and habitat-specialist species. The rapid pace of contemporary climate change often exceeds species’ adaptive capacity, resulting in demographic declines and range contractions (Breed *et al*., 2013; Shirey *et al*., 2024). Montane insects are particularly vulnerable, as their available habitat is inherently limited and may disappear entirely under future warming scenarios (Romo *et al*., 2023; Ursul *et al*., 2025).

Assessing the conservation status of insects remains challenging due to a lack of robust baseline data on population sizes, trends, and connectivity (IUCN, 2012). In this context, genetic information has become a key component of conservation biology (Garner *et al*., 2020). Small and isolated populations are especially susceptible to inbreeding and genetic drift, which erode genetic diversity and can trigger an “extinction vortex” (Frankham, 2005; Blomqvist *et al*., 2010; Palomares *et al*., 2012). Genetic analyses provide insights into population structure, levels of genetic diversity, inbreeding and effective population size, all of which are essential for informing conservation strategies and assessing extinction risk (Marí-Mena *et al*., 2019; Hohenlohe *et al*., 2021; González-Castellano *et al*., 2023; Sucháčková Bartoňová *et al*., 2023). Ecological Niche Modelling (ENM), or Species Distribution Modelling (SDM), is a powerful approach to predict species responses to climate change. By linking occurrence data to environmental variables, ENMs identify suitable habitats and project their spatial shifts under past, present or future climate scenarios (e.g., Sistri *et al*., 2022). For geographically restricted and ecologically specialized species, ENMs are particularly informative, revealing potential range contractions, altitudinal shifts, or poleward movements. For example, research on lowland butterflies has shown upward range shifts (Habel *et al*., 2023), a pattern that has also been particularly observed in mountain butterflies as species track cooler climates (Wilson *et al*., 2005; Rödder *et al*., 2021). Long-term monitoring reveals that mountain species with narrow climatic niches are especially vulnerable to warming, as suitable habitats become fragmented or disappear altogether. ENMs, therefore, provide essential tools for forecasting distributional range shifts and prioritising conservation efforts (Guisan & Thuiller, 2005; Settele *et al*., 2008). However, ENMs may underestimate extinction risk if species show local adaptation or if non-climatic constraints, such as host-plant availability, limit realised distributions (Harte *et al*., 2004).

The Spanish Moon Moth, *Actias isabellae* (Graells, 1849), is a flagship example of a montane specialist insect potentially threatened by global change. The species typically occurs at elevations between 700 and 1,700 meters, which correspond to its optimal ecological niche and provide climatic conditions suitable for its development. Marí-Mena et al. (Marí-Mena *et al*., 2016) detected pronounced phylogeographic structure in *A. isabellae*. Nuclear markers resolved six distinct genetic clusters corresponding to major mountain systems: French Alps, western and eastern Pyrenees, Central System, Iberian System, and Baetic Mountains. In contrast, mitochondrial variation showed a broader two-lineage pattern, grouping the Alps, Pyrenees, and Iberian System versus the Central System and Baetic Mountains. The traditionally recognized subspecies: *A. isabellae galliaegloria* (French Alps), *A. isabellae paradisea* (eastern Pyrenees), *A. isabellae roncalensis* (western Pyrenees), and *A. isabellae ceballosi* (Baetic range), occupy regions that match the nuclear clusters. Its striking morphology and restricted distribution have made *A. isabellae* an iconic European moth, but also a target for collectors (Rougeot, 1974). Intense collecting pressure during the 1970s led the mayors of five municipalities in the Queyras region to issue municipal decrees banning the collection of *A. isabellae galliaegloria*. These decrees were issued in 1975, one year before the adoption of the first French nature protection law. This subspecies is currently listed under Article 3 of the Decree of 23 April 2007 establishing the lists of insects protected throughout the French national territory and the conditions for their protection.

In French Alps, the subspecies *A. isabellae galliaegloria* occurs mainly in the departments of Alpes-de-Haute-Provence and Hautes-Alpes, with a higher concentration in the Queyras and the Guil Valley (Maurel *et al*., 2013; Breton *et al*., 2018). The origin of the Alpine subspecies has long been debated. Some authors considered a native origin unlikely and suggested a human-mediated introduction from Spain during the early 20th century (Chrétien, 1925; Bollow, 1932; Agenjo, 1943; Templado *et al*., 1975; Fernández-Vidal, 1992), whereas others argued for a native origin (Cleu, 1925, 1939; Warnecke, 1943; Maso-Planas & Ylla-Ullastre 1989; Nässig, 1991). More recent genetic analyses based on mitochondrial and microsatellite markers indicate that the Alpine subspecies is genetically unique and related to the eastern Iberian and Pyrenean populations (Marí-Mena *et al*., 2016). When it comes to foraging preferences, larvae of *A. isabellae galliaegloria* feed mainly on Scots pine (*Pinus sylvestris*), making the moth highly dependent on the distribution and health of its host plant. *A. isabellae galliaegloria* can be locally abundant in the Alps (Maurel *et al*., 2013) and may have benefited from extensive afforestation programs during the 19th century using *P. sylvestris* and *Pinus nigra*, which may have facilitated colonisation of new areas (Maurel *et al*., 2013; Leraut, 2020). But Scots pine is sensitive to drought and heat stress and has experienced stand-level dieback in southern and central Europe in recent decades (Bose *et al*., 2024). This raises concerns about the long-term viability of *A. isabellae galliaegloria* habitats under ongoing climate change, especially given that previous genetic studies revealed that the Alpine subspecies exhibits extremely low genetic diversity, compared to most Iberian populations (Marí-Mena *et al*., 2016). Despite this genetic impoverishment, these observations have led some authors to argue that the species should no longer be considered endangered in the Alpine region and that its protected status should be reconsidered (Leraut, 2020). Confirming the genetic status of the Alpine subspecies and understanding its demographic history and environmental constraints are, therefore, critical for assessing extinction risk and guiding conservation decisions.

In this study, we address two main objectives. First, we genetically characterise the French Alpine population of *A. isabellae galliaegloria* using RADseq (Restriction-site Associated DNA sequencing; (Baird *et al*., 2008). This genomic approach provides much higher resolution than previous studies based on mitochondrial markers and microsatellites (Marí-Mena *et al*., 2016), allowing us to investigate population structure, genetic diversity, and demographic history, and to assess the extent of genetic erosion. Second, we investigate the ecological constraints shaping the current and future distribution of the Alpine subspecies using species distribution models that incorporate both climatic variables and the distribution of its host plant, *P. sylvestris*. Using ENM, we project habitat suitability under current conditions and future climate scenarios (2050) to evaluate potential range shifts and risks of habitat contraction.

By coupling genomic data and ecological niche modelling, we aim to answer the following questions: (i) What is the current genetic status of the isolated Alpine subspecies of *A. isabellae*? (ii) How will climate change and host plant availability affect its future distribution and persistence in the Alps? Together, our results provide a robust framework for assessing the vulnerability of *A. isabellae galliaegloria* and for guiding conservation strategies for montane specialist insects facing rapid environmental change.

## Material and methods

### Sampling

In total 62 samples of *A. isabellae* collected during previous studies (Marí-Mena *et al*., 2016) were included in this study (Supplementary Table 1). These comprised 53 individuals of the Alpine subspecies *A. isabellae galliaegloria* from seven localities in the French Alps (Figure 1B), seven individuals of *A. isabellae paradisea* from two localities in the eastern Pyrenees and two individuals of *A. isabellae roncalensis* from one locality in the western Pyrenees (Supplementary Table 1). To capture the genetic diversity of the French Alpine subspecies as comprehensively as possible, sampling was focused on its main distribution areas in the administrative departments of Alpes-de-Haute-Provence and Hautes-Alpes, especially in the Guil Valley and Queyras massif (Figure 1B) under the appropriate collecting permits: Arrêtés n° 2008.38.4 and n° 2012-156-0008 issued by the Préfecture des Hautes-Alpes, Arrêté n° 2012.978 issued by the Préfecture des Alpes-de-Haute-Provence, permit ref LC/mp 24/2009/1638 issued by Gobierno de Aragón, and permit ref. 0152S issued by the Generalitat de Catalunya to CLV and MV. Sampling was non-lethal and consisted of removing a single leg or a piece of the hindwing tail following established protocols (Vila *et al*., 2009).

**Figure 1.**
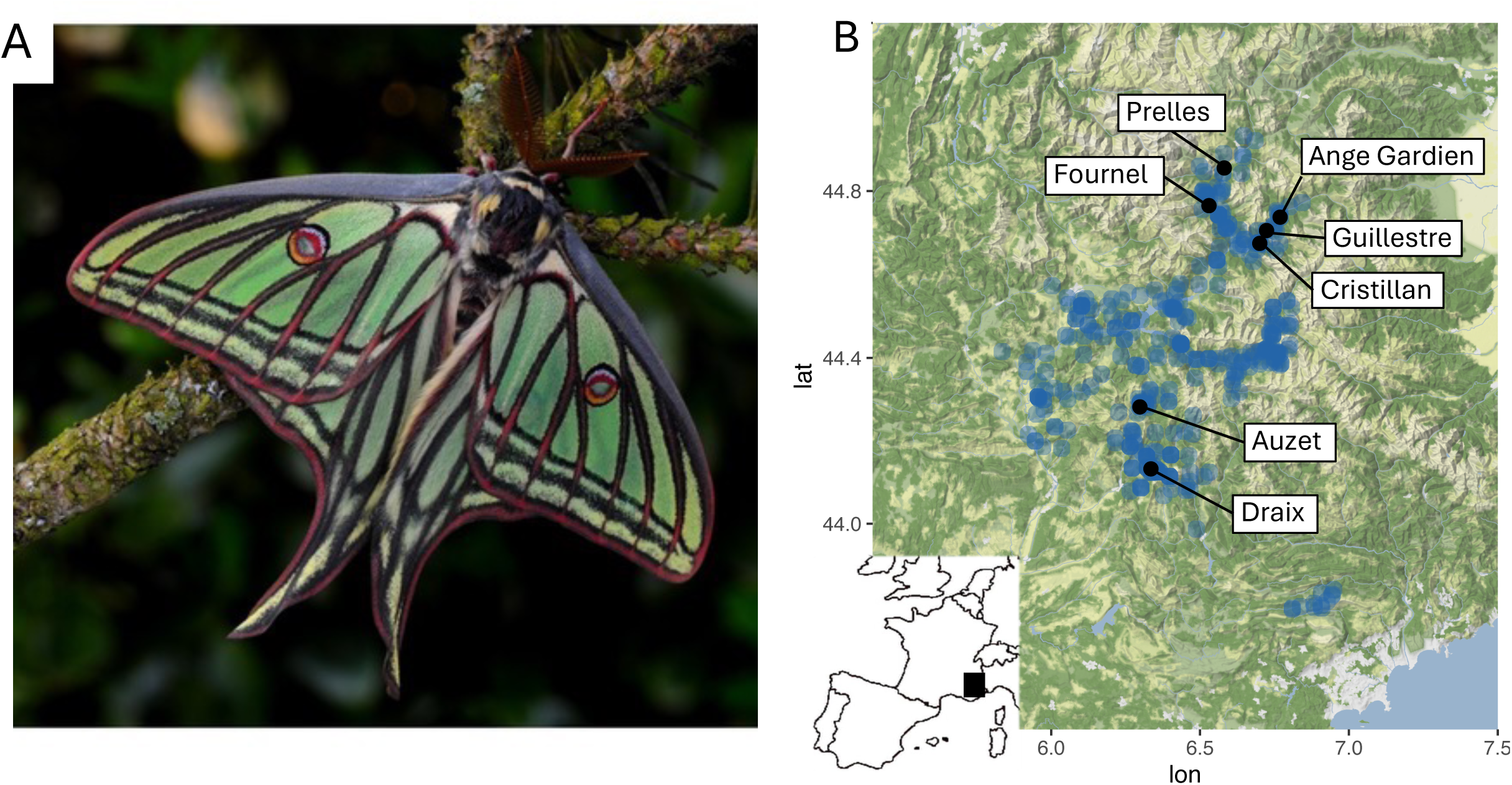
A. Habitus of a male of *A. isabellae galliaegloria* (credits: Thibaud Decaens). B. Map of the distribution in the Alps, the blue outline reflects the known distribution of *A. isabellae galliaegloria* using the GBIF occurrences (including the Gréolières locality in South of France) and black the localities included in the genetic analyses (n=7 in Prelles; n=9 in Fournel; n=9 in Ange Gardien; n=6 in Guillestre; n=7 in Cristillan, n=6 in Auzet and n=9 in Draix).

### DNA extraction, RADseq library and sequencing

DNA was extracted using the QIamp® mini kit from Qiagen, following the manufacturer’s instructions and protocol. Legs were cut in small pieces with a scalpel and incubated with the buffer and proteinase K in a ThermoMixer for at least 17 hours at 56°C and 400 rpm. The DNA was eluted in 50 μL of Nuclease-free water, with an incubation time of 30 minutes at room temperature before centrifugation. DNA concentrations were measured using the Qubit® 2.0 Fluorometer (ThermoFisher), using the dsDNA HS kit. DNA quality was also checked using the NanoDrop™ One spectrophotometer based on the 260/280 and 260/230 absorbance ratios. For the preparation of the RADseq library, genomic DNA was initially digested with the Sbfl restriction enzyme, followed by size selection to retain fragments ranging from 200 to 400 bp using a PippinPrep HT 1.5% (SageScience). These steps were undertaken to selectively capture homologous loci for each individual. The sequencing library was subsequently constructed by incorporating indexes and barcodes for individual identification, along with sequencing Illumina adapters. The sequencing was performed using 100 bp single reads from the Illumina Novaseq6000 technology. The RADseq library preparation and sequencing were performed by the MGX-Montpellier GenomiX (Montpellier, France).

### Population genetics analyses

The raw read quality was assessed using FastQC (https://www.bioinformatics.babraham.ac.uk/projects/fastqc/), demultiplexed and cleaned using process_radtags from *Stacks* (Rochette *et al*., 2019). Reads were mapped on the reference genome of *A. isabellae* genome (Vila *et al*., 2025) using *bwa mem* (Li, 2013). Mapping files were filtered according to the mapping quality, i.e. mapq < 20, using samtools (Li *et al*., 2009), and indels were realigned using the *Picard suite* (Picard toolkit, 2019). Variants were called using freebayes 1.3.8 (Garrison & Marth, 2012). Bi-allelic SNPs shared by at least 60% of the samples and with a minimum quality of 50 were extracted using vcftools (Danecek *et al*., 2011). Independent SNPs were extracted to perform genetic structure inference, keeping SNPs separated by at least 400 bp, thus corresponding to different RAD loci. To investigate genetic population structure, two different approaches were used. Population admixture was assessed using *STRUCTURE 2.3.3* (Pritchard *et al*., 2000) and testing K-value from 1 to 10. For each value, three runs were performed with a burn-in of 100,000 followed by 500,000 iterations. The Evanno method (Evanno *et al*., 2005) implemented in *Structure Harvester* (Earl & vonHoldt, 2012) was used to investigate the likelihood of the tested K-values. As a complementary approach, Principal Component Analysis (PCA) implemented for genetic data in the *adegenet* R package (Jombart & Ahmed, 2011) was used. Population differentiation was assessed using pairwise *F*ST (Weir & Cockerham, 1984) with the *pairwise.WCfst* function from the *hierfstat* R package (Goudet, 2005).

Genetic diversity indices, i.e. observed heterozygosity (*Ho*), expected heterozygosity (*He*) and the inbreeding coefficient (*F*IS) were estimated using the *basic.stats* function from the *hierfstat* R package (Goudet, 2005). The contemporary and historical effective population size (*N*e) were inferred using *GONE2* (Santiago *et al*., 2020) on the SNPs shared by all the samples. To optimise the number of samples that is too low at the locality scale, we combined all the samples from the Alps and performed a subsampling approach by resampling 30 individuals, 10 iterations, to test the robustness of the estimates. Genetic structure was accounted for using a model that considers a random set of individuals from a metapopulation composed of equally sized subpopulations, closely reflecting the studied system. A constant recombination rate of 3 cM/Mb across the genome was applied (Yamamoto *et al*., 2008).

### Occurrences data

At the french national scale, occurrence data for *A. isabellae galliaegloria* were primarily obtained through a monitoring program documenting its distribution in the French Alps (Maurel *et al*., 2013; Breton *et al*., 2018). This program, initiated in 2012 and coordinated by INRAE Orléans in collaboration with the naturalist association Proserpine, was conducted during the emergence period in May and June using a synthetic sex pheromone (Millar *et al*., 2010). It enabled the identification of species presence points across the Alpine range. These data were combined with records from previous studies in the French Alps and Pyrenees (Millar *et al*., 2010; Vila *et al*., 2010; Lopez-Vaamonde & Goussard, 2011; Maurel *et al*., 2013; Morichon *et al*., 2014; Marí-Mena *et al*., 2016, 2019; Breton *et al*., 2018)

Additional occurrence records were obtained from publicly available databases, including Artemisiae (https://oreina.org/artemisiae/index.php) and the Global Biodiversity Information Facility (GBIF;https://doi.org/10.15468/dl.u54nhc; Flemons *et al*., 2007). To ensure data quality, points with imprecise coordinates (<1000m), duplicates, erroneous localities, and erratic male records were removed. The French dataset contained 655 occurrences, resulting in 370 unique records after filtering. To reduce spatial autocorrelation and improve analytical robustness, the dataset was thinned to 262 occurrences using the *thin* function from the *spThin* package (Aiello-Lammens *et al*., 2015), selecting one occurrence per raster cell (∼1 km²) using randomization and iteration approaches. For the modelling approach (see below), 250 presence records were retained from the initial set of 262, after excluding the putative introduced population as explained in the result section, i.e. Gréolières locality in southern France (Figure 1). For the Principal Component Analysis (PCA), an additional dataset composed of 186 Spanish occurrences was obtained from GBIF and filtered using the same criteria applied to the French dataset, resulting in a global dataset of 446 occurrences.

### Environmental variables

To predict the potential current and future distribution of *A. isabellae galliaegloria* using Ecological Niche Modelling (ENM), we downloaded the 19 bioclimatic variables for the current period (1981–2010) and future projections for 2050 from the CHELSA database (https://chelsa-climate.org/; Brun *et al*., 2022) at a spatial resolution of 30 arc-seconds (∼1 km²). From the 19 bioclimatic variables, BIO8, BIO9, BIO18 and BIO19 were excluded due to spatial artefacts (Campbell *et al*., 2015), resulting in 15 remaining variables. Subsequently, a subset of these variables was transformed following CHELSA guidelines (scale and offset) to ensure that raster values matched their corresponding units of measurement. First, we performed Principal Component Analysis (PCA) including these 15 variables using the *ade4* R package (Dray *et al*., 2007) to compare the climatic niches of the Spanish and Alpine populations. To carry out the PCA, bioclimatic values were extracted at each occurrence point using the *RChelsa* package (Karger *et al*., 2017), and normalized.

To further perform the analyses at the study scale, we cropped the downloaded layers at the following spatial extent: longitude 4.960 to 7.186 and latitude 43 to 47.575. A dendrogram was then constructed using the *hclust* function from the R *stats* package. To reduce multicollinearity, variables with a pairwise Pearson correlation coefficient < 0.7 were selected for ENM construction under current and future conditions. Three bioclimatic variables were retained: BIO1 (Annual Mean Temperature), BIO3 (Isothermality), and BIO15 (Precipitation Seasonality). Future climate projections were based on the IPCC Shared Socioeconomic Pathways (ssp1-2.6, ssp3-7.0, ssp5-8.5), representing different socioeconomic trajectories, for the years 2050 and 2080, separately. Projections were obtained using the MPI-ESM1-2-HR model with CMIP6 simulations (Müller *et al*., 2018).

Current and future distributions of *P. sylvestris*, and the primary host plant of *A. isabellae galliaegloria*, were also included. The current distribution was obtained from land-use data in BD Forêt® V2, available via the Institut national de l’information géographique et forestière geoservice (IGN, https://geoservices.ign.fr/bdforet). For future projections, we used a 2050 distribution under the rcp8.5 scenario provided by ClimEssences (Piboule *et al*., 2021). To ensure consistency with CHELSA layers, the ClimEssences projection was resampled to a spatial resolution of 30 arc-seconds.

### Modelling approach

Species distribution models at the Alps scale were developed using the three selected bioclimatic variables and the *P. sylvestris* distribution, and projections were generated using the R package *biomod2* (Guéguen *et al*., 2025). To account for algorithmic variability in species distribution projections (Thuiller *et al*., 2019), we used an ensemble modelling framework (Araújo & New, 2007) incorporating eight statistical algorithms: Maximum Entropy (MaxEnt; Phillips *et al*., 2006), Random Forest (RF; Breiman, 2001), Random Forest downsampled (RFd), Flexible Discriminant Analysis (FDA; Hastie *et al*., 1994), Generalized Linear Model (GLM; Nelder & Wedderburn, 1972), Generalized Boosting Model (GBM; Friedman, 2001), Generalised Additive Model (GAM; Hastie & Tibshirani, 1986), and Extreme Gradient Boosting (XGBoost; Chen & Guestrin, 2016).

As these algorithms require either presence–absence data or presence-only data supplemented with pseudo-absences, and true absence data were unavailable, we first generated pseudo-absence points using a random sampling strategy with 10 replicates of 750 points each. Indeed, we generated three times more pseudo-absences than presence data based on advice from the *biomod2* authors. Model validation was performed using a random cross-validation strategy with five repetitions, allocating 80% of the data for training and 20% for testing. Equal weighting (0.5) was applied to presences and pseudo-absences, and variable importance was assessed through 10 permutations.

A total of 400 single models were generated under current environmental conditions for each of two sets of variables: one using only climatic variables (first model), and the other combining climatic variables with *P. sylvestris* distribution (second model). Models were evaluated using the Boyce index (Hirzel *et al*., 2006), which is specifically suited for assessing predictive performance of presence-only models. This index is a continuous metric, with theoretical values ranging from -1 (model predicts the opposite of the observed occurrences) to 1 (perfect agreement between model predictions and observed occurrences), and with 0 corresponding to a performance no better than random. An ensemble modelling approach was then applied to combine individual models, retaining only those with a Boyce index above 0.7. Variable importance was estimated for the ensemble model using a permutation procedure repeated over 10 runs. Final ensemble models were constructed using three methods: weighted mean (EMwmean), coefficient of variation (EMcv), and committee averaging (EMca). Models with Boyce values > 0.7 were retained for the final ensemble forecasts. Independent evaluation based on the Boyce index showed better performance of the EMwmean compared to the EMca. Consequently, only EWmean is presented in the remainder of the results, and was used to project suitability and prediction presence. We defined areas of high habitat suitability by applying a conservative threshold of 0.6 to the EMwmean ensemble projections, retaining only pixels predicted as highly suitable across models. Habitat suitability was projected for present-day and future (2050) climatic scenarios. We measured spatial overlap between the predicted suitability (first vs. second model and current vs. future scenarios) using the *dismo* R package (Hijmans *et al*., 2017) by calculating Schoener’s D and Warren’s I overlap metrics (Warren *et al*., 2008). Those metrics range from 0 to 1, with values close to 1 indicating a larger overlap. In total, two current projections, with and without *P. sylvestris* distribution, and four future projections, under three climatic scenarios and including *P. sylvestris* for the corresponding one, ssp5-8.5, were generated.

## Results

### Genetic characterisation of A. isabellae galliaegloria

The RADseq protocol and subsequent sequencing resulted in a total of 46 million reads, i.e. an average of 0.75 million reads per sample (sd = 0.18 million reads) (Supplementary Table 1). The mapping of the reads on the reference genome revealed a high percentage of mapping, an average of 96.4%. The variant calling resulted in a total of 1.2 million raw genetic variations. The filtering retained 5,007 shared SNPs suitable for performing population genomic estimates and characterising the state of the populations. Only one of the initial 62 samples was discarded due to an excessive proportion of missing data, up to 20% of missing.

Genetic structure analyses first revealed a strong differentiation between the Alpine subspecies, *A. isabellae galliaegloria* and the two Pyrenean subspecies *A. isabellae paradisea* and *A. isabellae roncalensis.* This separation is evident in the admixture analysis, with the most likely number of clusters K = 2 (Figure 2B, Supplementary Table 2), and in the PCA along PC1, which explains 31.07% of the genetic variance and clearly separates samples from the two mountain ranges (Figure 2C). High levels of genetic differentiation are also reflected in *F*ST values, which exceed 0.5 for each pair of genetic clusters from the Alps versus the Pyrenees (Figure 3).

**Figure 2.**
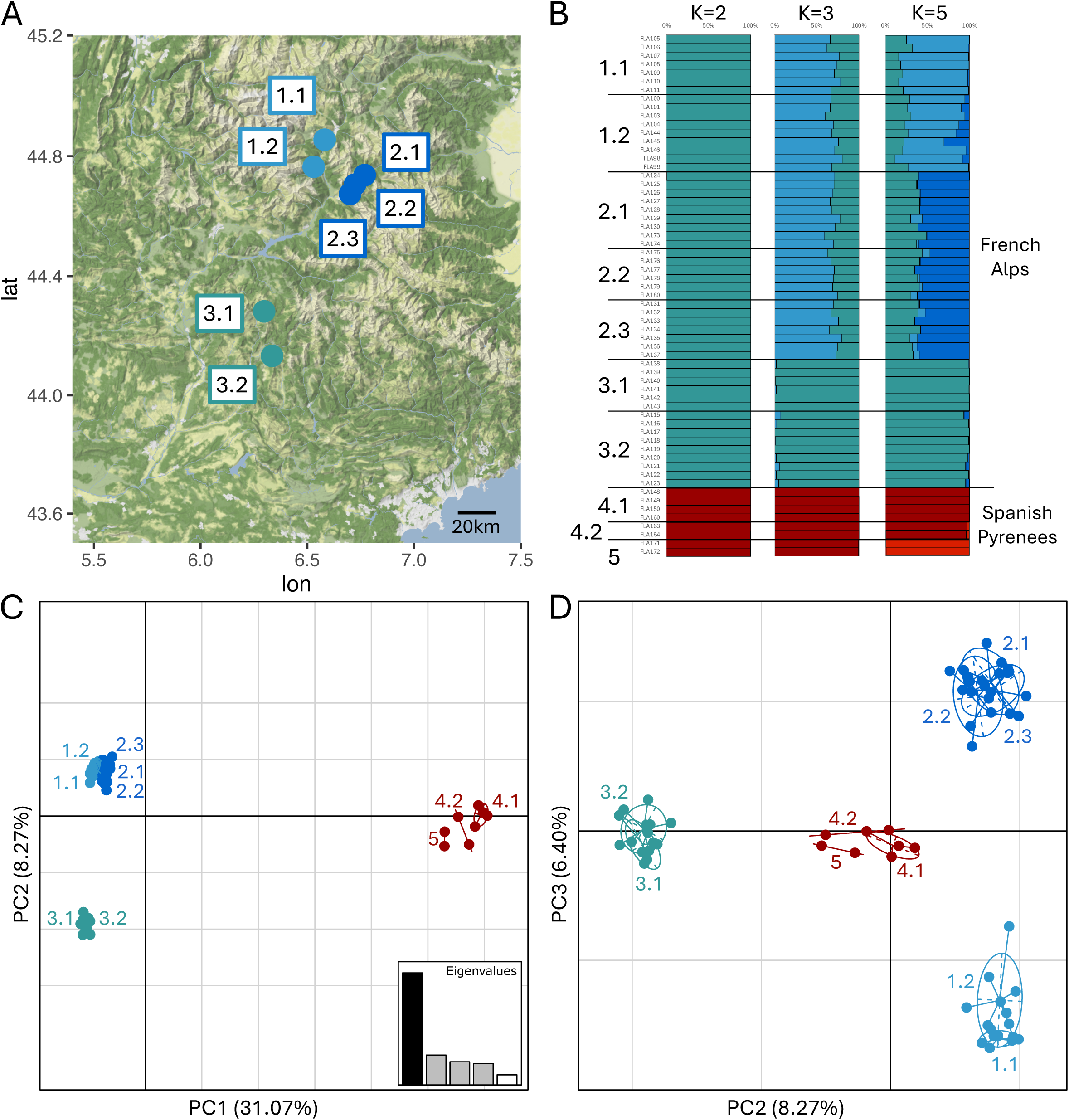
Genetic structure of French populations of *A.isabellae*. A. Geographic distribution of the three genetic clusters, 1.1 Prelles; 1.2 Fournal; 2.1 Ange Gardien; 2.2 Guillestre; 2.3 Cristillan, 3.1 Auzet, 3.2 Draix. B. Barplot of the individual admixture levels for K=2, K=3 and K=5. C. D. Principal component analysis representing the sample distribution along the first PCs, respectively on PC1/PC2 and PC2/PC3. The Alpine genetic cluster is depicted in blue tones, and the western and Eastern Pyrenean clusters in red.

**Figure 3.**
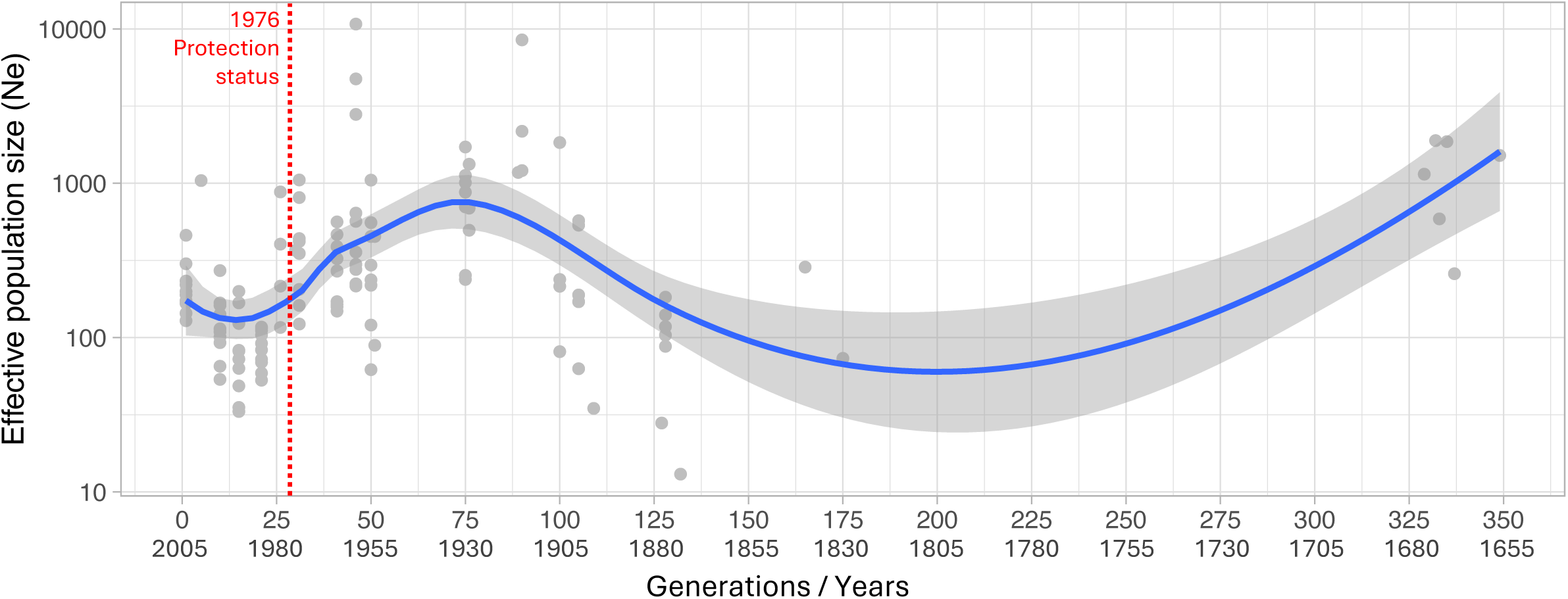
Effective population size (*N*e) over time since 350 years ago. Effective population size is plotted from the present to the past. The blue line represents the median effective population size. The shaded area represents the 95% confidence interval (CI 95%).

Within the Alpine range, a substructure among localities was detected. This substructure appears in both the admixture results at K = 5 (Figure 2B) and in the PCA, which identifies three genetic clusters (Figure 2 C & D). In the PCA, PC2 (8.27% of variance) separates cluster 3 (Figure 2D), which includes the southern Alpine localities of Auzet and Draix (Figure 1C), while PC3 (6.40% of variance) separates the two northern Alpine clusters along an east–west gradient (Figure 1D). *F*ST values among these three Alpine clusters are lower, ranging from 0.182 to 0.296, than between the Alpine and Spanish populations (Supplementary Figure 2). Although the western Pyrenean subspecies *A. isabellae roncalensis* was represented by a single locality (Gillué) with only two individuals sequenced, the K = 5 analyses (Figure 2B) revealed a clear split from the eastern Pyrenean subspecies *A. isabellae paradisea* (seven specimens from two localities, Baiasca and Embouni) (Supplementary Figure 1 and Supplementary Table 1), with an *F*ST value of 0.227. This level of genetic differentiation between the western (*roncalensis*) and eastern (*paradisea*) Pyrenean lineages is comparable to that observed among the three Alpine genetic clusters. However, admixture proportions among the Pyrenean populations were more mixed, possibly reflecting ongoing gene flow or incomplete lineage sorting resulting from recent divergence.

Genetic diversity was assessed using observed heterozygosity (*Ho*) and mean gene diversity within populations (*Hs*) at both the locality and genetic cluster levels. *Ho* and *Hs* estimates were consistent among localities within the same Alpine cluster, and cluster-level estimates obtained by pooling localities yielded similar values (Table 1), indicating that relatively small sample sizes did not bias the results.

**Table 1.** Descriptive statistics (genetic cluster, locality, number of samples, *Ho*= observed heterozygosity, *Hs*= mean gene diversity within populations, *F*IS= inbreeding coefficient).

| Population | | Locality | N | $H_o$ (se) | $H_s$ (se) | $F_{IS}$ (se) |
| --- | --- | --- | --- | --- | --- | --- |
| 1 | 1,1 | Prelles | 7 | 0.046 (0.002) | 0.055 (0.002) | 0.104 (0.018) |
|  | 1,2 | Fournel | 9 | 0.053 (0.002) | 0.062 (0.002) | 0.127 (0.017) |
|  |  | merged | 16 | 0.050 (0.002) | 0.061 (0.002) | 0.189 (0.015) |
| 2 | 2,1 | Ange Gardien | 9 | 0.064 (0.002) | 0.074 (0.002) | 0.132 (0.015) |
|  | 2,2 | Guillestre | 6 | 0.067 (0.002) | 0.076 (0.002) | 0.088 (0.016) |
|  | 2,3 | Cristillan | 7 | 0.063 (0.002) | 0.073 (0.002) | 0.104 (0.015) |
|  |  | merged | 22 | 0.065 (0.002) | 0.075 (0.002) | 0.192 (0.013) |
| 3 | 3,1 | Auzet | 6 | 0.039 (0.002) | 0.047 (0.002) | 0.130 (0.022) |
|  | 3,2 | Draix | 9 | 0.040 (0.002) | 0.050 (0.002) | 0.178 (0.019) |
|  |  | merged | 15 | 0.039 (0.002) | 0.049 (0.002) | 0.234 (0.018) |
| 4 | 4,1 | Baixasca | 4 | 0.168 (0.003) | 0.190 (0.003) | 0.049 (0.010) |
|  | 4,2 | Embonui | 2 | 0.192 (0.004) | 0.213 (0.004) | -0.082 (0.014) |
|  |  | merged | 6 | 0.174 (0.003) | 0.202 (0.003) | 0.086 (0.008) |
| 5 |  | Gillue | 2 | 0.200 (0.004) | 0.228 (0.004) | -0.035 (0.013) |

Among Alpine clusters, heterozygosity varied from *Ho* = 0.039 in the least diverse cluster (Auzet) to *Ho* = 0.067 in the most diverse cluster (Guillestre). Compared to the Pyrenean localities, Alpine populations exhibited substantially lower genetic diversity, up to fivefold lower in Auzet versus Embonui (Table 1). A similar pattern was observed for the inbreeding coefficient (*F*IS), which was significantly higher in Alpine localities, whereas Pyrenean populations showed values near zero, consistent with an absence of inbreeding. When samples were pooled at the cluster level, *F*IS increased, potentially reflecting a Wahlund effect, where population subdivision leads to an apparent deficit of heterozygotes relative to Hardy–Weinberg expectations (Hartl & Clark, 1997) (Table 1).

The historical trajectory of the effective population size (*N*e) between 1655 and 1900 is associated with greater uncertainty but suggests a gradual decline, potentially linked to extensive deforestation in the Alpine region. This was followed by a subsequent increase in *N*e, reaching a maximum around 1930, likely due to widespread afforestation programs (Figure 3). The population then gradually declined until the mid-1990s, possibly as a result of intensive collecting pressure. There are indications of a renewed increase in *N*e since that period, which may reflect the establishment of legal protection for the subspecies in 1976.

### Niche characterisation

Thanks to the rich occurrence dataset, from both the Alpine populations and the Spanish range where the species is widely distributed (Figure 4A), we were able to encompass the full environmental breath of the species. As an exploratory step, we first performed a PCA on all environmental variables extracted for the 446 unique occurrences (Figure 4B). The first two principal components accounted for 92% of the variance in the dataset clear separation between Alpine and Spanish occurrences is visible along PC1, with only limited overlap. These results suggest that the Spanish and Alpine populations occupy contrasted ecological niches.

**Figure 4.**
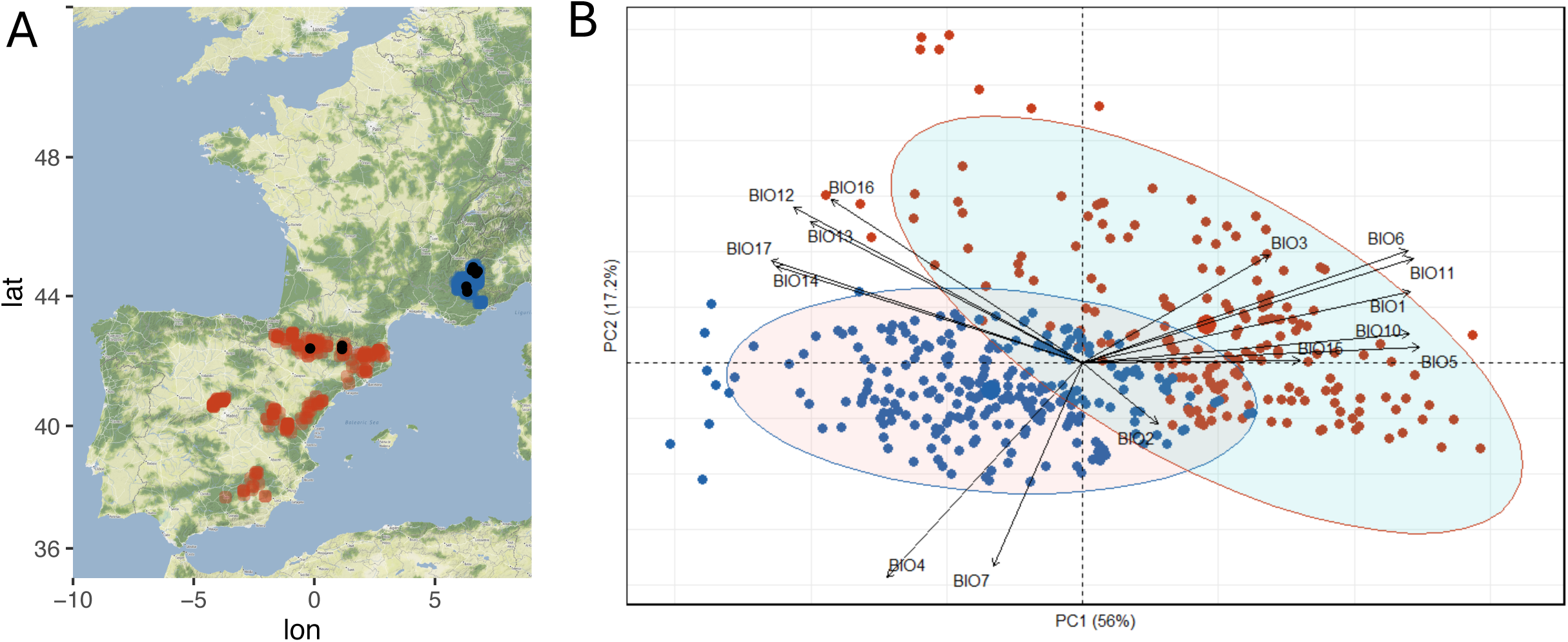
A. Map of the occurrence distribution in Spain (in red) and Alps (in blue). Black dots indicate the locality sampled for the genetic analyses. **B** Climatic PCA with the 15 bioclimatic variables from Spanish (red) and French (blue) occurrences.

It should be noted that an additional locality in southern France (Maritime Alps), Gréolières, where individuals of *A. isabellae* have recently been recorded (unpublished data from C. Lopez-Vaamonde) was initially included in the Alpine dataset. However, this locality displayed an unexpected position in the PCA, clustering instead with Spanish populations (Supplementary Figure 3). Furthermore, there is a possibility that individuals may have been deliberately released by amateur breeders, which is consistent with its distinct ecological conditions. Because of this uncertainty, and to avoid biasing the niche modelling of the Alpine population, these occurrences were removed from the final dataset.

To precisely characterise the potential distribution of the Alpine population and its future evolution, an Ecological Niche Modelling (ENM) approach was implemented. For both modelling approaches (climate-only and climate + *P. sylvestris*), all single models except the Generalised Additive Model (GAM) produced high predictive accuracies, as indicated by Boyce Index values ranging from 0.77 to 0.97 with an average of 0.91 and were incorporated into the ensemble modelling framework (Supplementary Figure 4, Supplementary Table 3). The GAM algorithm, with a Boyce index of 0.43 for the climate-only modelling approach, fell below the selection threshold of 0.7 and was therefore excluded from the ensemble. The strong Boyce values indicate robust model training and a high ability to discriminate suitable from unsuitable habitats under the current climatic conditions. Ensemble models (EMwmean, EMca, and EMcv) integrated all high-performing single models, yielding spatially coherent predictions of suitable habitat for *A. isabellae* under current conditions. The convergence across ensemble metrics indicates low model uncertainty and high agreement among algorithms (Supplementary Figure 4).

### *Drivers of* A. isabellae galliaegloria *distribution*

Across algorithms, the relative contributions of the environmental predictors were consistent, suggesting stable model behaviour despite methodological differences among algorithms. For both modelling approaches, climatic predictors related to precipitation patterns were the dominant determinants of *A. isabellae* distribution. In particular, precipitation seasonality (BIO15) emerged as the most influential factor (accounting for >80 % of total model contribution for climate-only), indicating that variation in rainfall across seasons strongly constrains habitat suitability for the species in the Alps. In contrast, temperature-related variables (BIO1 and BIO3) contributed less to model performance. Indeed, BIO1 exhibited a moderate contribution (20%), whereas BIO3 contributed only marginally (<1%), implying that the current range is more tightly associated with moisture regimes than with thermal conditions. When *P. sylvestris* distribution - which covers 13,36 % of the 14,471 km^2^ study area-was incorporated into the modelling framework, the explanatory power of precipitation variables remained dominant (>50%), while the Scots pine variable showed a secondary contribution (10%), suggesting the species’ dependence on its host plant. This confirms that both climatic suitability and host plant availability jointly determine the realised niche of *A. isabellae*.

Under current climatic conditions, ensemble models consistently identified suitable habitats for *A. isabellae* throughout the region surrounding its current distribution area, namely the westernmost part of the Alps, and supporting the robustness of the ensemble predictions (Figure 1). The most favourable area is the meridional Pre-Alps, west of the Mercantour massif, which has more moderate altitudes (Figure 5). Further to the north-east, a dendritic pattern can be observed in the highest massifs, such as the Ecrins and Queyras, following the valleys and revealing that the favourable habitat does not cover the highest altitudes. There is also an extension of favourable habitat towards the north-west, reaching almost as far as the Vaucluse massif, even though the species has never been observed there. Low suitability values were observed both further north and further south. In the north, the predicted range shows a clear limit around 45°N, even in areas that might appear favourable, such as the Vanoise massif. In the south, including the Mercantour region, lower suitability likely reflects the stronger Mediterranean climatic influence. Across the entire study area (Figure 5), which covers a total of 128,160 km², the habitat area considered favourable, with a suitability >0.6, covers 9,765 km², or 7.62%. When incorporating the distribution of *P. sylvestris*, the extent of suitable habitat was spatially congruent with the climate-only projections (Figure 5). In the same study area, excluding the edges in Italy and Switzerland due to the lack of data for the Scots pine distribution, the total area considered covers 99,777 km², of which 7,640 km² are considered favourable, and representing 7.66% of the study area. This indicates that host-plant availability acts as an additional constraint refining the climatic niche envelope of *A. isabellae*. The overlap between both modelling approaches (climate-only vs. climate + *P. sylvestris*) was high (Schoener’s D = 0.81; Warren’s I = 0.93), suggesting that bioclimatic and biotic drivers operate synergistically to determine the realized distribution of the species. These results should be considered with caution, as the distribution of *P. sylvestris* used in the modelling framework was itself derived from a species distribution model based on occurrence data, but also on the same set of abiotic predictors.

**Figure 5.**
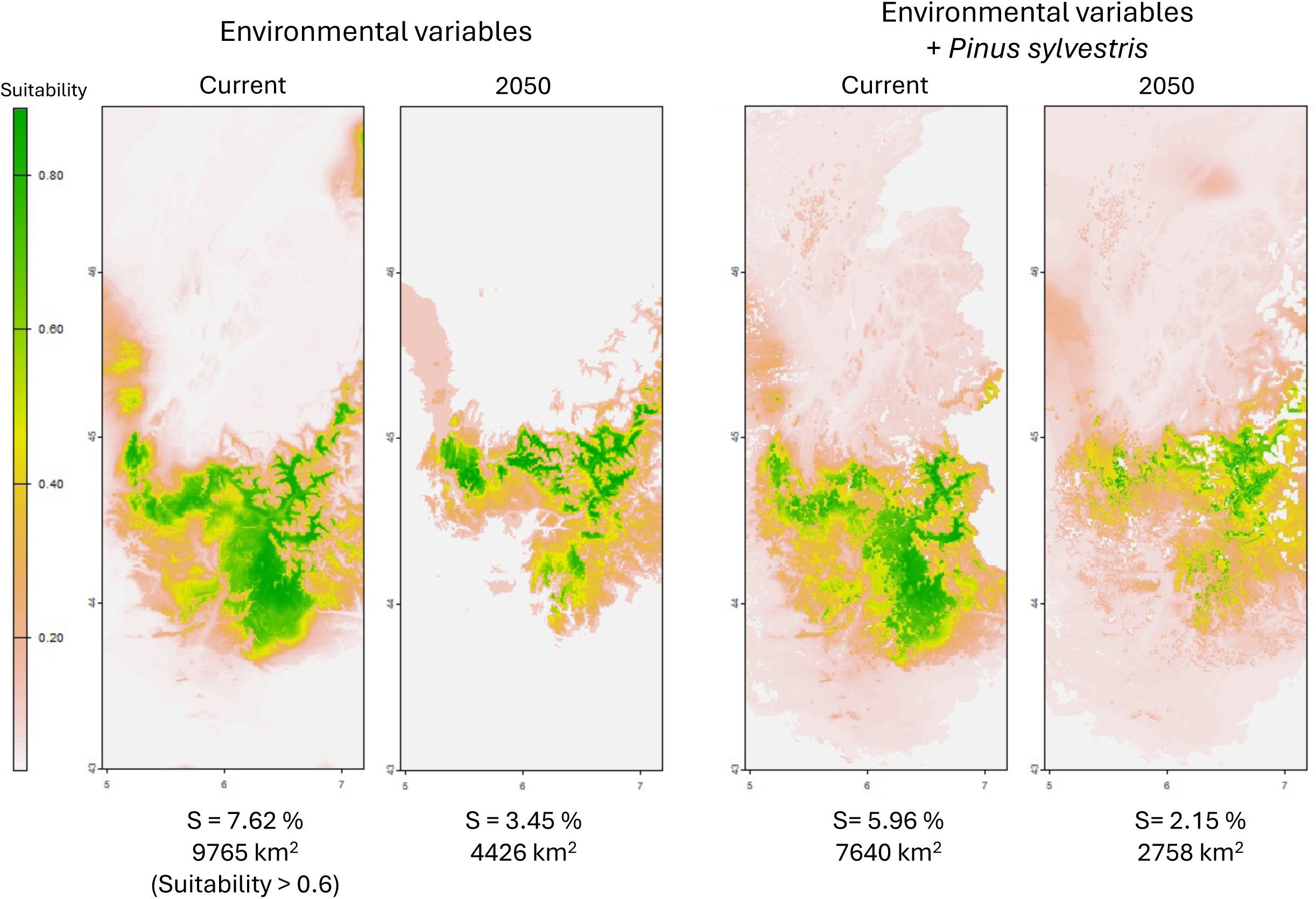
Species distribution models of *A. isabellae* under current and future (2050) conditions, based on bioclimatic variables (BIO1, BIO3 and BIO15) and Scots pine distribution. Suitability values range from low (grey) to high (green). Percentages indicate the proportion of the study area with suitability >0.6 and the corresponding surface for each scenario.

The mean overlap between current and future distribution models was also high for the climate-only (Schoener’s D = 0.76; Warren’s I = 0.95) and for the climate + *P. sylvestris* (Schoener’s D = 0.66; Warren’s I = 0.86) approach, suggesting strong similarity in suitability patterns between current and future projections.

### Prediction of suitable areas under future climatic conditions

The comparison of predicted suitable habitats for the current and the future (2050) reveals a contraction in suitable habitat area for the three ssp scenarios tested, with variations in the intensity of this reduction (Figure 5). This difference in intensity is rather substantial, ranging from an 11% loss of suitable habitat (1,085 km²) with 8,680 km² of remaining favourable habitat for the most optimistic scenario (ssp1-2.6) to a loss of nearly 55% (5,339 km²) for the most pessimistic scenario (ssp5-8.5), resulting in a suitable habitat area of 4,426 km². The intermediate scenario (ssp3-7.0), also shows an intermediate loss of 36% of favourable habitat area (3,524 km²). In the three scenarios, the pattern of loss is consistent, with losses mainly occurring in the southern part of the distribution area (Figure 5). However, there is no counterbalancing effect towards the north with a putative favourable habitat extension above the 45°N limit. This artificial limit is still marked in the three inferences (Figure 5). Similarly, there does not appear to be any altitudinal extension of the distribution area, or if there is, it does not increase the total area of suitable habitat. The integration of Scots pine distribution projections for 2050 into the inference and under the most pessimistic scenario (ssp5-8.5 for abiotic data and corresponding to the prediction of *P. sylvestris* under rcp8.5) reveals an even more drastic contraction in suitable habitat area. This area is reduced by 64% to 2,758 km². These results reveal that *P. sylvestris* appears to be even more sensitive to future climate change. That said, even though *A. isabellae* appears to be less sensitive; its intrinsic biological dependence on Scots pine makes it no less vulnerable.

## Discussion

### Current situation

The analysis of population structure and the pronounced genetic differentiation between localities in the French Alps and the Pyrenees supporting the native origin of the Alpine lineage and refuting the hypothesis of a human-mediated introduction of Iberian individuals in the early 20th century (Chrétien, 1925; Agenjo, 1943). The observed genetic structure is consistent with previous microsatellite-based results (Marí-Mena *et al*., 2016), which clearly separated the two Pyrenean subspecies. However, the higher resolution of the RADseq dataset (5,007 SNPs vs. nine microsatellite loci) reveals finer-scale structure, identifying three distinct genetic clusters within the Alps. These clusters show a clear geographic structuring and indicate reduced gene flow across the mountain ranges. The fine-scale genetic substructure shown here for the Alpine subspecies must be integrated into conservation strategies, as it directly influences the spatial distribution and long-term maintenance of genetic diversity.

Similar topographic constraints on dispersal have been reported in *Lycaena* species (Trense *et al*., 2022), where the more sedentary *L. hippothoe* exhibited, in the Alps, strong population structure and localized genetic diversity, consistent with low dispersal ability and valley isolation. Landscape-scale genetic structuring shaped by topography and land-use barriers has also been documented in *Phengaris* butterflies (Gauthier *et al*., 2025). Although *A. isabellae* has a large wingspan and potentially higher dispersal capacity, female sedentary behaviour likely limits effective dispersal. Males actively search for pheromone-emitting females at the end of the flight period, when calling females are scarce, and occasionally disperse away from core areas (Maurel *et al*., 2013), with vagrant males observed in adjacent regions such as l’Oisans (Baillet, 2008). Despite male mobility, female behaviour largely defines species distribution, as shown in *Phengaris* (Plazio *et al*., 2020), where females preferred valley corridors and avoided crossing hills, unlike more mobile males.

The comparison of genetic diversity between Alpine and Pyrenean populations reveals substantially lower diversity in the Alps. These estimates are directly comparable, as they were obtained using the same methodology and a similar number of nuclear loci, providing robust insight into population status. They are consistent with previous microsatellite-based findings (Marí-Mena *et al*., 2016), with observed heterozygosity (*H*_O_) two to three times higher in Pyrenean populations. They are also aligned with the low diversity observed at the mitochondrial level, with a single haplotype, which more specifically reflects the evolutionary trajectory of females. This reduced diversity in the Alps could be the result of a founder effect during the colonization of the Alps, but this process seems to have taken place a long time ago, and diversity should have emerged in a healthy population (Dapporto *et al*., 2024). The most likely mechanism linked to this low diversity would be recent effective population size variations, and more specifically declines and bottlenecks. The isolation of the Alpine population seems to have then led to inbreeding and amplify the effects of genetic drift, pushing progressively this population into a vortex of extinction (Bosse & van Loon, 2022).

Although cross-study comparisons of diversity metrics are often complicated by differing methodologies and species-specific life histories, the heterozygosity values observed in Alpine *A. isabellae* rank among the lowest reported for European butterflies (Després *et al*., 2019; Trense *et al*., 2021; Kebaïli *et al*., 2023; Gauthier *et al*., 2025), including species considered threatened (Dupuis *et al*., 2020) or critically endangered (Nakahama *et al*., 2024). Low genetic diversity constrains adaptive potential by limiting the allelic pool available for natural selection, thereby weakening responses to pathogens, climatic changes, or habitat shifts. It also increases susceptibility to inbreeding depression, as reflected in elevated inbreeding coefficients. Isolation combined with reduced diversity raises the likelihood of matings between related individuals, increasing the risk of homozygosity for deleterious alleles, which typically manifests as reduced fitness through lower survival, fertility, and developmental stability (Dussex *et al*., 2023). In conservation contexts, such patterns are widely recognized as warning signals of limited evolutionary resilience and heightened extinction risk (Nakahama *et al*., 2024).

Effective population size (*N*e) is a key indicator of genetic health and accurately reflecting genetic dynamics (Waples, 2025). In some Spanish *A. isabellae* populations, despite apparently healthy census sizes (*N*c), genetic monitoring has revealed strikingly low *N*e values. For instance, in one Pyrenean locality, *Ne* was estimated at only 27–49, despite ca. 1500 reported individuals (Marí-Mena *et al*., 2019). This *Ne*/*Nc* ratio implies that the population genetic effective size is much smaller than its census size, consistent with unequal reproductive success and/or a reduced effective number of breeders. This increases the rates of genetic drift and inbreeding and can accelerate the loss of neutral genetic diversity.

Historical demographic inference suggests a rise in *N*e between 1800 and 1930, potentially linked to extensive afforestation programs in the Alps using both *P. sylvestris* and *P. nigra laricio* (Leraut, 2020). Alpine reforestation effectively began with the French law of 28 July 1860, which established both voluntary (subsidized) and compulsory (including possible expropriation) reforestation zones, followed by rapid implementation from 1861 onward with large designated areas, nurseries, and seed facilities (France, Law of 28 July 1860; Fourchy, 1963). In Alpes-de-Haute-Provence alone, 500–1,000 ha were reforested annually between 1895 and 1914 (Douguedroit, 1980). The more recent trajectory, from the beginning of the 20th century, appears to be multifactorial linked to deforestation, contemporary forest management as well as the gradual influence of global warming, that have reduced the size and connectivity of pine stands, resulting in habitat fragmentation. Smaller and isolated patches increase the risk of local extinctions and impede recolonization, reinforcing the evolutionary isolation of populations.

The collection pressures have certainly also played a role in the recent decline. Indeed, as an emblematic species, *A. isabellae* has been widely collected by amateur entomologists throughout its range and in particularly in the Alps (Rougeot, 1974), prompting local declines and the implementation of protective measures. By the mid-20th century, conservation concerns led to full legal protection in several countries, including France in 1976, where it became the first insect to receive such status.

The benefits of this protection may already be observable, since the 1990s, *Ne* has increased slightly. However, short-term demographic gains may mask ongoing genetic erosion due to the lag between demographic recovery and genetic response. Genetic diversity does not collapse immediately after a population decline; rather, losses accumulate gradually over successive generations. This phenomenon, known as genetic extinction debt or drift debt, refers to genetic variation that is effectively doomed to disappear. Simulations and empirical studies (Liu *et al*., 2025) show that small populations continue to lose diversity long after a bottleneck, even if census sizes remain stable, due to the cumulative effects of drift and inbreeding. This process likely affects Alpine populations of *A. isabellae galliaegloria,* as evidenced by their low diversity and elevated inbreeding. Once lost, recovery of genetic variation is slow. Maintaining adaptive potential thus depends not only on current diversity but also on preserving sufficiently large and connected populations to ensure gene flow and minimise inbreeding. Long-term conservation will require the maintenance of favourable conditions to support these processes.

### Putative evolution of the Alpine subspecies

The long-term persistence of Alpine populations of *A. isabellae galliaegloria* depends critically on maintaining sufficient genetic diversity, effective population sizes, and habitat connectivity. However, these prerequisites appear increasingly challenged under projected future environmental conditions as revealed by this work. Indeed, our ENMs indicate a contraction of suitable habitat, largely driven by climate change and the patchy distribution of its primary host plant, *P. sylvestris*. Among the environmental variables tested, annual mean temperature and precipitation seasonality explained most of the variation in the current distribution of the Alpine subspecies. These factors collectively shape the species’ ecological niche, confining it to mid- to high-elevation montane habitats where microclimatic conditions and host plant availability coincide.

Genetic structuring in mountainous species typically reflects historical isolation in glacial refugia, limited dispersal and topographic barriers (Trense *et al*., 2021). The genetic structure of the Alpine populations of *A. isabellae* further illustrates the influence of landscape configuration on evolutionary trajectories. Three distinct genetic clusters were identified, corresponding broadly to the northern, central, and southern sectors of the French Alps. This substructure reflects the role of valleys and mountain ridges as barriers to gene flow, restricting dispersal and reinforcing genetic differentiation. Such isolation, combined with small effective population sizes, may exacerbate the loss of allelic diversity over time, limiting adaptive potential and increasing vulnerability to stochastic events. Landscape connectivity is therefore a critical determinant of both genetic and demographic stability in these montane populations. ENMs also suggest that future climatic conditions will further constrain the distribution of *A. isabellae*, with a pronounced poleward and upward shift in suitable habitats. In addition to this latitudinal migration, a significant contraction of the overall range is predicted, particularly in lower-elevation valleys where warming and drought reduce host plant availability. Comparisons with other European montane insects indicate that upward or poleward shifts are common responses to warming, though local microclimatic refugia can buffer populations against broader trends (Biella *et al*., 2024). While ENMs provide a useful forecast, fine-scale topographic variation may create pockets of suitable habitat that enable local persistence despite overall range contraction (Stark & Fridley, 2022).

The distribution of *A. isabellae* is tightly constrained by the availability and continuity of *P. sylvestris*, its obligate larval host. The past demographic trajectory appears to follow the forest dynamics of *P. sylvestris* as mentioned above. But an additional factor potentially influencing the future trajectory of Alpine *A. isabellae* is the capacity for host-plant flexibility. While larvae predominantly feed on *P. sylvestris*, the species can also utilize other pine species (Maso-Planas & Ylla-Ullastre 1989; Nässig, 1991; Gázquez *et al*., 2008; Sánchez-Fernández & de Arce-Crespo, 2017). In the context of climate warming, a host shift toward higher-altitude *Pinus uncinata* or more drought-tolerant *P. pinaster* could provide opportunities for range persistence or even expansion into new habitats. Such flexibility may partially buffer populations against local declines in *P. sylvestris* due to drought, heat stress, or forestry practices.

Taken together, these results suggest that the evolutionary trajectory of Alpine *A. isabellae galliaegloria* populations is shaped by a combination of historical demographic events, landscape configuration, and host plant availability. While the species exhibits some capacity for dispersal, especially in males, landscape barriers and habitat fragmentation impose limits that may constrain future adaptive responses (González-Castellano *et al*., 2023). Conservation efforts should therefore focus not only on preserving existing populations but also on maintaining or restoring habitat connectivity, ensuring the persistence of *P. sylvestris* patches, and monitoring microclimatic refugia that may buffer populations against climate-driven range contractions. This integrated approach is essential to mitigate the compounded risks of genetic erosion and habitat loss and to promote long-term evolutionary resilience in this emblematic montane insect.

## Data Accessibility

Raw genomic data can be found under NCBI BioProjects PRJNAXXXXX. VCF files and custom scripts have been made available in the Github repository (https://github.com/JeremyLGauthier/Actias_isabellae_Conservation_genomics). The GBIF occurrence datasets are available at https://doi.org/10.15468/dl.u54nhc and https://doi.org/10.15468/dl.rk6bcs.

## Acknowledgements

We thank François Breton for providing occurrence data. We are also grateful to François-Xavier Saintongue (DSF, URZF) for his valuable comments on modelling the distribution of *Pinus sylvestris* in France using ClimEssences.

