## Supplementary Material for "Genetic erosion and projected habitat loss in the protected Alpine moth *Actias isabellae galliaegloria* (Lepidoptera: Saturniidae)"

**Supplementary Table 1.** Descriptive statistics comprising sampling information (localities, coordinates), molecular statistics (concentration, purity), and sequencing.

| Code | Country | Locality | Coordinates | C (ng/µl) | Ratio 280/260 | Ratio 260/230 | # reads | % mapping | # shared SNPs |
|---|---|---|---|---|---|---|---|---|---|
| FLA100 | France | Fournel | 44°45′55.3″N 6°31′51.4″E | 32.8 | 2.1 | 2.07 | 1045449 | 94.48 | 4857 |
| FLA101 | France | Fournel | 44°45′55.3″N 6°31′51.4″E | 23.7 | 2.13 | 1.7 | 674887 | 97.34 | 4817 |
| FLA103 | France | Fournel | 44°45′55.3″N 6°31′51.4″E | 32.8 | 2.13 | 2.18 | 747285 | 97.8 | 4831 |
| FLA104 | France | Fournel | 44°45′55.3″N 6°31′51.4″E | 23.4 | 2.06 | 1.67 | 715713 | 96.42 | 4808 |
| FLA105 | France | Prelles | 44°51'17.6"N 6°34'52.6"E | 26.4 | 2.14 | 1.65 | 660034 | 97.73 | 4769 |
| FLA106 | France | Prelles | 44°51'17.6"N 6°34'52.6"E | 22 | 2.15 | 1.62 | 816848 | 97.27 | 4823 |
| FLA107 | France | Prelles | 44°51'17.6"N 6°34'52.6"E | 29.4 | 2.14 | 1.59 | 722204 | 97.18 | 4843 |
| FLA108 | France | Prelles | 44°51'17.6"N 6°34'52.6"E | 22.1 | 2.05 | 1.53 | 892698 | 94.56 | 4891 |
| FLA109 | France | Prelles | 44°51'17.6"N 6°34'52.6"E | 31.3 | 2.12 | 2.08 | 722445 | 98.07 | 4801 |
| FLA110 | France | Prelles | 44°51'17.6"N 6°34'52.6"E | 26.8 | 2.14 | 2.08 | 776044 | 98.06 | 4812 |
| FLA111 | France | Prelles | 44°51'17.6"N 6°34'52.6"E | 18.8 | 2.24 | 1.56 | 786511 | 97.18 | 4842 |
| FLA115 | France | Draix | 44° 7'57.41"N 6°20'6.09"E | 23.7 | 2.11 | 1.65 | 667364 | 95.49 | 4825 |
| FLA116 | France | Draix | 44° 7'57.41"N 6°20'6.09"E | 25.3 | 2.09 | 1.62 | 552797 | 97.17 | 4753 |
| FLA117 | France | Draix | 44° 7'57.41"N 6°20'6.09"E | 24.4 | 2.13 | 1.6 | 690886 | 94.08 | 4823 |
| FLA118 | France | Draix | 44° 7'57.41"N 6°20'6.09"E | 20.7 | 2.15 | 1.54 | 585477 | 95.2 | 4771 |
| FLA119 | France | Draix | 44° 7'57.41"N 6°20'6.09"E | 28 | 2.13 | 2.04 | 646825 | 93.8 | 4808 |
| FLA120 | France | Draix | 44° 7'57.41"N 6°20'6.09"E | 20.5 | 2.2 | 1.4 | 304994 | 97.4 | 4694 |
| FLA121 | France | Draix | 44° 7'57.41"N 6°20'6.09"E | 25.2 | 2.09 | 1.37 | 846119 | 97.67 | 4853 |
| FLA122 | France | Draix | 44° 7'57.41"N 6°20'6.09"E | 22 | 2.05 | 1.63 | 698215 | 98.33 | 4820 |
| FLA123 | France | Draix | 44° 7'57.41"N 6°20'6.09"E | 11.9 | 2.12 | 1.48 | 1049763 | 98.11 | 4832 |
| FLA124 | France | Ange_Gardien | 44°44′14.6″N 6°46′8.31″E | 13.9 | 2.14 | 1.59 | 667636 | 98.06 | 4865 |
| FLA125 | France | Ange_Gardien | 44°44′14.6″N 6°46′8.31″E | 15 | 2.18 | 1.5 | 763928 | 97.11 | 4854 |
| FLA126 | France | Ange_Gardien | 44°44′14.6″N 6°46′8.31″E | 15.3 | 2.16 | 1.69 | 563899 | 98.19 | 4818 |
| FLA127 | France | Ange_Gardien | 44°44′14.6″N 6°46′8.31″E | 17.4 | 2.15 | 1.86 | 640202 | 97.71 | 4826 |
| FLA128 | France | Ange_Gardien | 44°44′14.6″N 6°46′8.31″E | 13.8 | 2.18 | 1.56 | 526707 | 98.02 | 4812 |
| FLA129 | France | Ange_Gardien | 44°44′14.6″N 6°46′8.31″E | 15.8 | 2.13 | 1.57 | 547433 | 98.01 | 4795 |
| FLA130 | France | Ange_Gardien | 44°44′14.6″N 6°46′8.31″E | 12.4 | 2.16 | 1.42 | 548781 | 97.64 | 4766 |
| FLA131 | France | Cristillan | 44°40′29.0″N 6°42′02.2″E | 12.2 | 2.07 | 1.65 | 650595 | 97.05 | 4851 |
| FLA132 | France | Cristillan | 44°40′29.0″N 6°42′02.2″E | 14.7 | 2.08 | 1.54 | 742834 | 96.14 | 4842 |
| FLA133 | France | Cristillan | 44°40′29.0″N 6°42′02.2″E | 12.4 | 2.09 | 1.36 | 434829 | 97.85 | 4786 |
| FLA134 | France | Cristillan | 44°40′29.0″N 6°42′02.2″E | 17.2 | 2.09 | 1.78 | 707996 | 94.86 | 4859 |
| FLA135 | France | Cristillan | 44°40′29.0″N 6°42′02.2″E | 15.6 | 2.1 | 1.61 | 774043 | 97.12 | 4857 |

| | | | | | | | | | |
|---|---|---|---|---|---|---|---|---|---|
| FLA136 | France | Cristillan | 44°40’29.0”N 6°42’02.2”E | 19.2 | 2.19 | 1.56 | 971598 | 96.9 | 4872 |
| FLA137 | France | Cristillan | 44°40’29.0”N 6°42’02.2”E | 20.2 | 2.09 | 1.79 | 804994 | 97.46 | 4847 |
| FLA138 | France | Auzet | 44°16’55.6”N 6°17’52.6”E | 34.5 | 2.05 | 1.87 | 767431 | 98.16 | 4866 |
| FLA139 | France | Auzet | 44°16’55.6”N 6°17’52.6”E | 30.8 | 2.14 | 2.05 | 1078931 | 98.62 | 4859 |
| FLA140 | France | Auzet | 44°16’55.6”N 6°17’52.6”E | 28.1 | 2.09 | 1.8 | 668784 | 94.23 | 4809 |
| FLA141 | France | Auzet | 44°16’55.6”N 6°17’52.6”E | 32.3 | 2.11 | 2.15 | 941197 | 94.86 | 4862 |
| FLA142 | France | Auzet | 44°16’55.6”N 6°17’52.6”E | 28.5 | 2.1 | 2.04 | 736333 | 92.8 | 4836 |
| FLA143 | France | Auzet | 44°16’55.6”N 6°17’52.6”E | 33.5 | 2.06 | 1.81 | 758906 | 95.62 | 4845 |
| FLA144 | France | Fournel | 44°45’55.3”N 6°31’51.4”E | 34.1 | 2.13 | 1.94 | 795653 | 98.09 | 4821 |
| FLA145 | France | Fournel | 44°45’55.3”N 6°31’51.4”E | 29.9 | 2.08 | 1.94 | 821833 | 94.89 | 4872 |
| FLA146 | France | Fournel | 44°45’55.3”N 6°31’51.4”E | 28.1 | 2.1 | 1.95 | 898678 | 97.63 | 4831 |
| FLA148 | Spain | Baiasca | 42°30’07.2”N 1°08’59.0”E | 9.1 | 1.98 | 1.35 | 952977 | 95.07 | 4676 |
| FLA149 | Spain | Baiasca | 42°30’07.2”N 1°08’59.0”E | 16.1 | 2.03 | 1.77 | 756338 | 95.82 | 4690 |
| FLA150 | Spain | Baiasca | 42°30’07.2”N 1°08’59.0”E | 17.7 | 2.05 | 1.89 | 720755 | 94.11 | 4627 |
| FLA160 | Spain | Baiasca | 42°30’07.2”N 1°08’59.0”E | 12.6 | 2.05 | 1.57 | 1041547 | 94.64 | 4723 |
| FLA161 | Spain | Embonui | 42°23'52.9"N 1°08'52.8"E | 9.82 | 1.92 | 0.98 | 202872 | 93.82 | 3569 |
| FLA163 | Spain | Embonui | 42°23'52.9"N 1°08'52.8"E | 10.1 | 1.87 | 1.27 | 358652 | 95.74 | 4563 |
| FLA164 | Spain | Embonui | 42°23'52.9"N 1°08'52.8"E | 9.18 | 2.05 | 1.29 | 789723 | 93.01 | 4732 |
| FLA171 | Spain | Gillue | 42°25'06.5"N 0°10'24.3"W | 17.6 | 2.02 | 1.41 | 854514 | 96.39 | 4725 |
| FLA172 | Spain | Gillue | 42°25'06.5"N 0°10'24.3"W | 26.4 | 2.01 | 1.74 | 616673 | 97.39 | 4683 |
| FLA173 | France | Ange_Gardien | 44°44’14.6”N 6°46’8.31”E | 21.2 | 2.13 | 1.67 | 655396 | 97.86 | 4839 |
| FLA174 | France | Ange_Gardien | 44°44’14.6”N 6°46’8.31”E | 28.4 | 2.11 | 1.87 | 693294 | 97.91 | 4839 |
| FLA175 | France | Guillestre | 44°42’19.8”N 6°43’24.2”E | 27.5 | 2.03 | 1.8 | 1073291 | 97.68 | 4894 |
| FLA176 | France | Guillestre | 44°42’19.8”N 6°43’24.2”E | 32.6 | 2.13 | 2.16 | 995177 | 97.85 | 4885 |
| FLA177 | France | Guillestre | 44°42’19.8”N 6°43’24.2”E | 24.2 | 2.06 | 1.73 | 839904 | 97.2 | 4858 |
| FLA178 | France | Guillestre | 44°42’19.8”N 6°43’24.2”E | 25.5 | 2.08 | 1.79 | 870137 | 96.52 | 4888 |
| FLA179 | France | Guillestre | 44°42’19.8”N 6°43’24.2”E | 30.7 | 2.05 | 1.81 | 873696 | 97.37 | 4884 |
| FLA180 | France | Guillestre | 44°42’19.8”N 6°43’24.2”E | 33.2 | 2.09 | 2.16 | 857652 | 95.85 | 4843 |
| FLA98 | France | Fournel | 44°45’55.3”N 6°31’51.4”E | 26.6 | 2.13 | 1.56 | 887421 | 97.06 | 4826 |
| FLA99 | France | Fournel | 44°45’55.3”N 6°31’51.4”E | 31.2 | 2.13 | 2.14 | 889552 | 94.18 | 4792 |

**Supplementary Table 2.** STRUCTURE Likelihood

| # K | Reps | Mean LnP(K) | Stdev LnP(K) | Ln'(K) | \|Ln''(K)\| | Delta K |
|---|---|---|---|---|---|---|
| 1 | 3 | -43033.5000 | 0.8660 | NA | NA | NA |
| 2 | 3 | -30760.2333 | 1.4978 | 12273.266667 | 11036.866667 | 7368.836004 |
| 3 | 3 | -29523.8333 | 452.4699 | 1236.400000 | 7093.066667 | 15.676327 |
| 4 | 3 | -35380.5000 | 2660.8070 | -5856.666667 | 11208.833333 | 4.212569 |
| 5 | 3 | -30028.3333 | 2182.7839 | 5352.166667 | 11011.266667 | 5.044598 |
| 6 | 3 | -35687.4333 | 5313.4711 | -5659.100000 | 6906.733333 | 1.299853 |
| 7 | 3 | -34439.8000 | 3146.4113 | 1247.633333 | 3880.700000 | 1.233373 |
| 8 | 3 | -37072.8667 | 9348.8713 | -2633.066667 | 8543.933333 | 0.913900 |
| 9 | 3 | -31162.0000 | 2146.2320 | 5910.866667 | 6938.666667 | 3.232953 |
| 10 | 3 | -32189.8000 | 1427.9690 | -1027.800000 | 4699.100000 | 3.290758 |
| 11 | 3 | -37916.7000 | 1841.7041 | -5726.900000 | NA | NA |

**Supplementary Table 3.** BOYCE statistics for each model used in the niche modeling approach.

| Algorithms | Bioclimatic model | | Bioclimatic + Pinus model | |
|---|---|---|---|---|
| | BOYCE (Train) | BOYCE (Test) | BOYCE (Train) | BOYCE (Test) |
| Extreme Gradient Boosting | 0.77696 | 0.68268 | 0.86356 | 0.71362 |
| Generalised Additive Model | 0.43052 | 0.37142 | 0.73280 | 0.75238 |
| Generalised Boosting Model | 0.93960 | 0.74382 | 0.94026 | 0.79366 |
| Generalized Linear Model | 0.90092 | 0.82364 | 0.91704 | 0.83734 |
| Flexible Discriminant Analysis | 0.90400 | 0.79796 | 0.87016 | 0.72868 |
| Random Forest downsampled | 0.94410 | 0.79182 | 0.95568 | 0.86068 |
| Random Forest | 0.94856 | 0.79246 | 0.97320 | 0.84410 |
| Maximum Entropy | 0.97356 | 0.79246 | 0.96944 | 0.88796 |

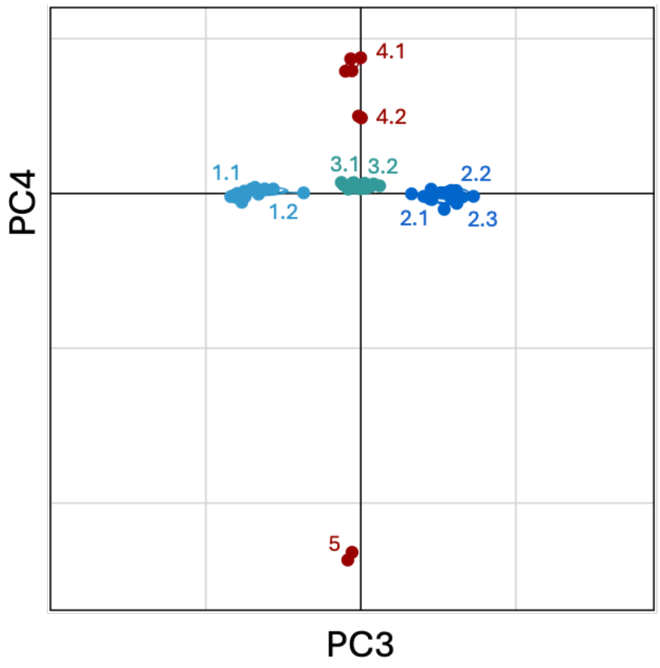


**Supplementary Figure 1.** Principal component analysis representing the sample distribution along the PC3 and PC4. The Alpine genetic cluster is depicted in blue tones, and the western and eastern Pyrenean clusters in red.

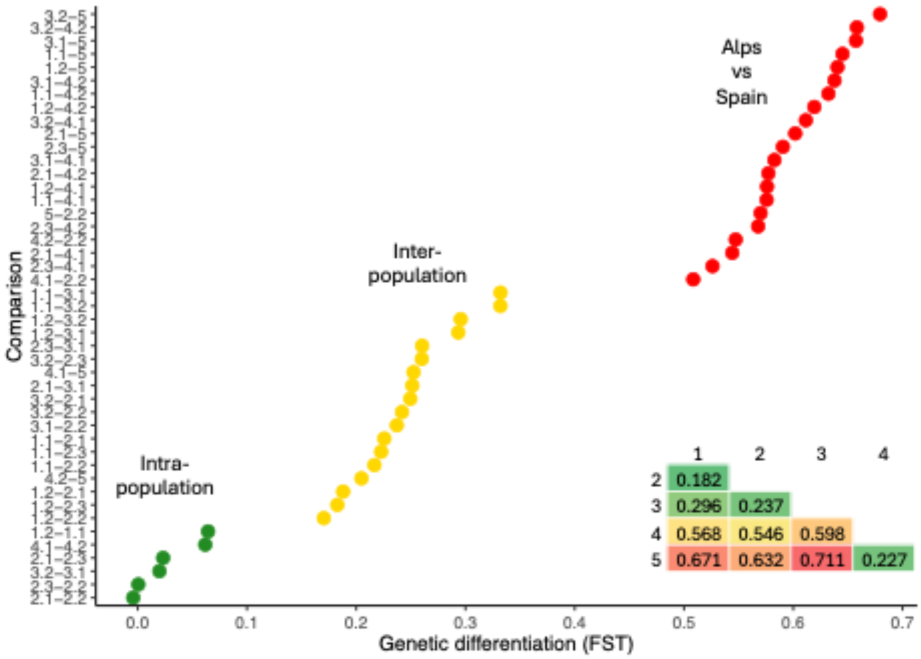


**Supplementary Figure 2**. $F_{ST}$ between localities and populations

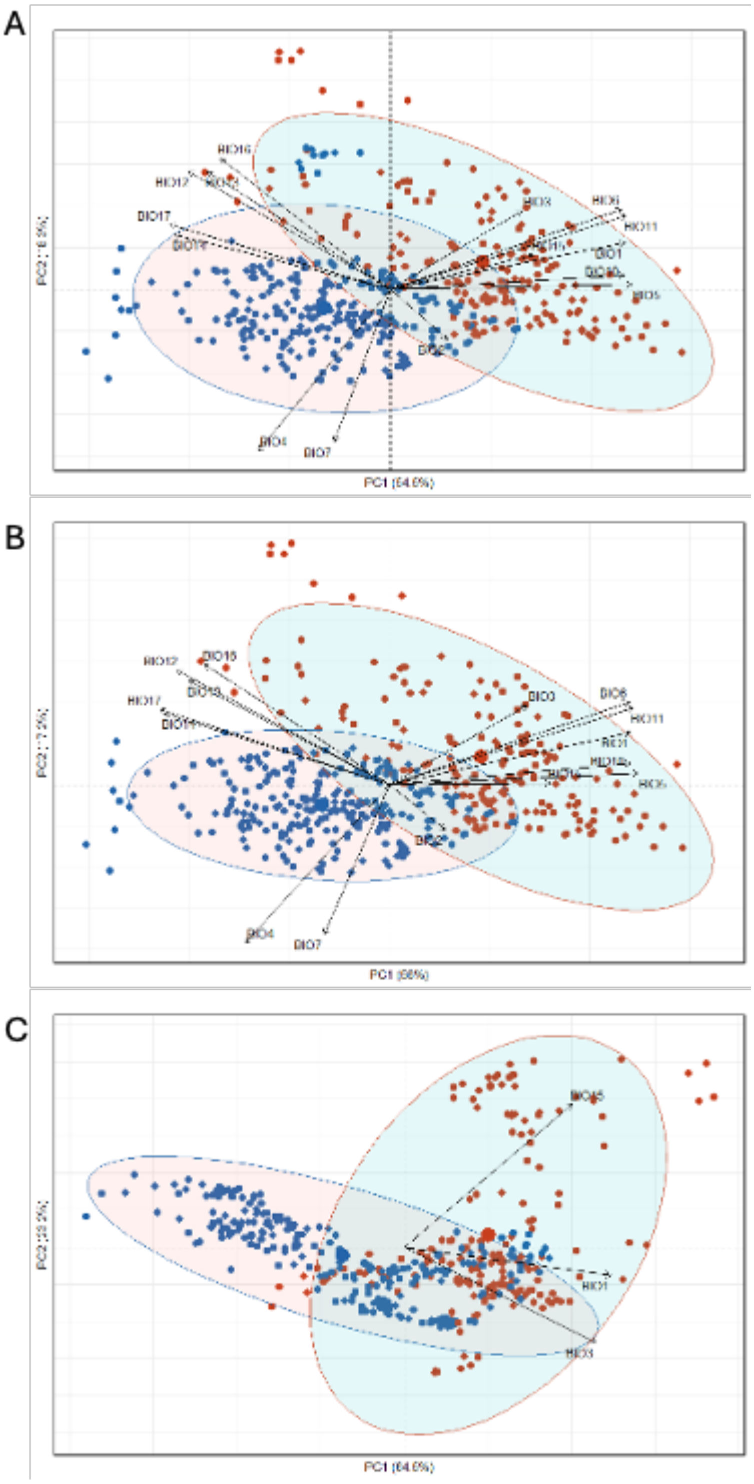


**Supplementary Figure 3.** A PCA including all samples (from Spain, Alps and Gréolières locality) and using all 19 environmental variables B PCA including all "natural" samples (without Gréolières locality) and using all 19 environmental variables C PCA including all "natural" samples (without Gréolières locality) and using the three selected environmental variables (BIO1, BIO3, and BIO15).

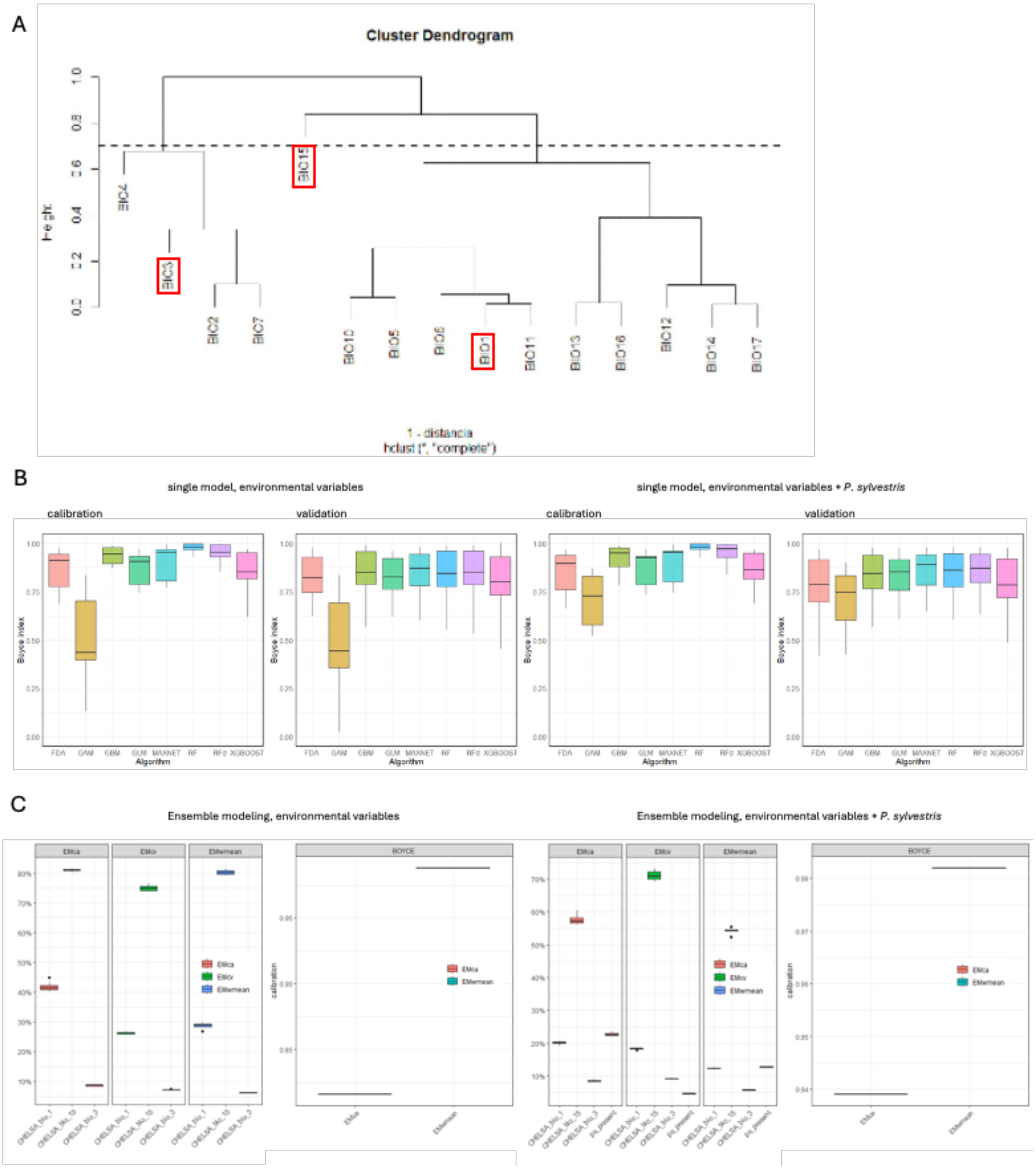


**Supplementary Figure 4.** Ensemble model construction and performance assessment of single models.

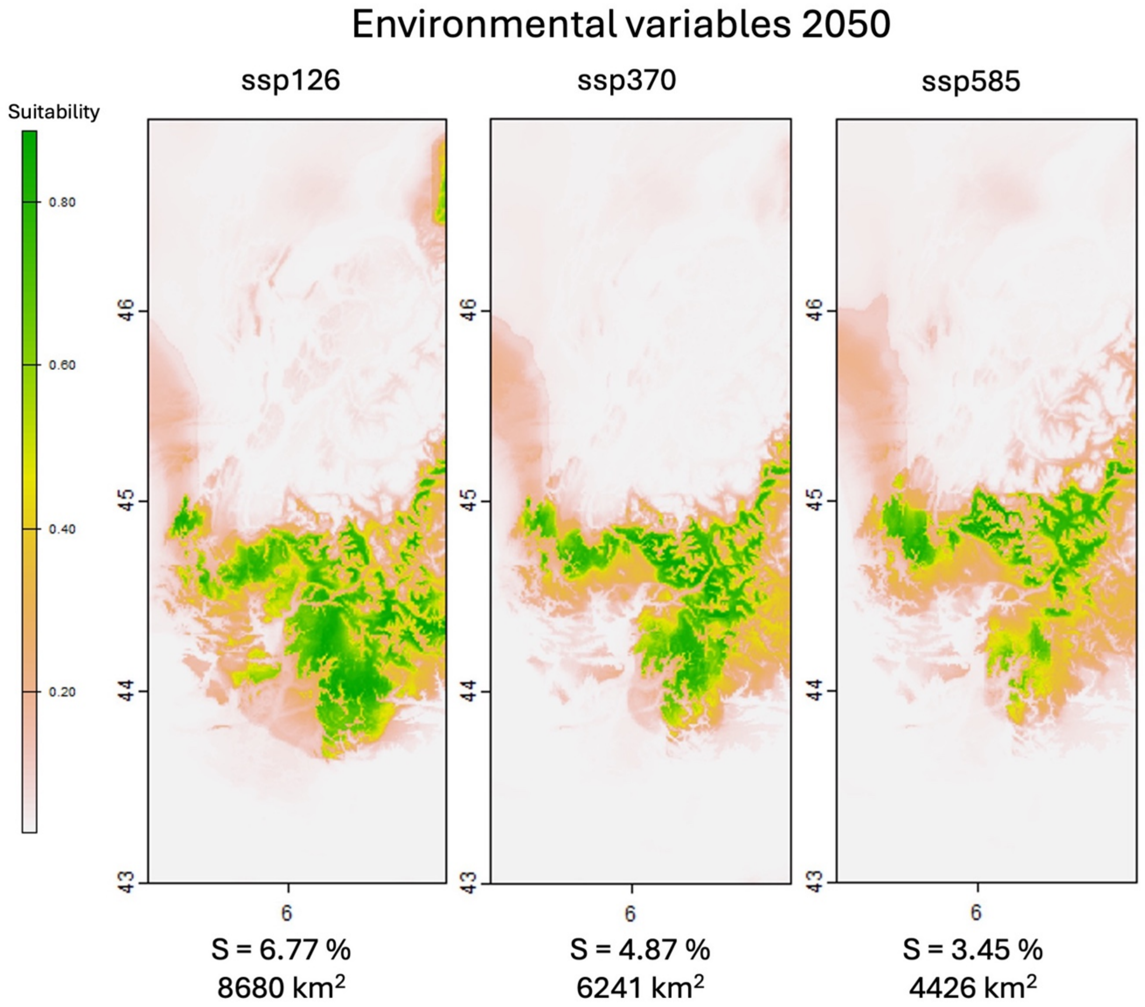


**Supplementary Figure 5.** Species distribution models of *A. isabellae* under future (2050) conditions, based on bioclimatic variables (BIO1, BIO3 and BIO15) and three climatic scenarios, ssp126, ssp370, and ssp585. Suitability values range from low (grey) to high (green). Percentages indicate the proportion of the study area with suitability >0.6 and the corresponding surface for each scenario.
